# Caloric Restriction Promotes Ischemia/Reperfusion Cardioprotection Through Increased Mitochondrial Na^+^/Ca^2+^ Exchange

**DOI:** 10.64898/2026.07.20.739602

**Authors:** Maiara I.C. Queiroz, Camille C. Caldeira da Silva, Éverton Vogt, Marcos A.E. Cruz, Julian D.C. Serna, Luiz R.G. Bechara, Julio C.B. Ferreira, Heberty T. Facundo, Alicia J. Kowaltowski

**Author notes:** Correspondence: Alicia J. Kowaltowski - Av. Prof. Lineu Prestes, 748, Cidade Universitária, 05508-000, São Paulo, SP, Brazil.

## Abstract

Caloric restriction (CR) protects against cardiac ischemia/reperfusion (I/R) injury, but the underlying mechanisms remain incompletely understood. Since mitochondrial Ca^2+^ overload and redox imbalance is a major driver of cardiac damage during reperfusion, we investigated if enhanced mitochondrial Ca^2+^ efflux contributes toward CR-induced cardioprotection. Rats were subjected to 16 weeks of *ad libitum* (AL) feeding or 40% caloric restriction. CR significantly increased mitochondrial Ca^2+^ retention capacity when Na^+^ ions were present, reduced H_2_O_2_ release, and increased the expression of proteins related to mitochondrial Ca^2+^ extrusion. In cardiomyocytes exposed to serum from CR rats, Ca^2+^ retention capacity also markedly increased, as well as Ca^2+^ efflux and the expression of extrusion proteins NCLX and TMEM65. Following I/R, CR hearts exhibited improved functional recovery accompanied by enhanced retention and increased mitochondrial Ca^2+^ efflux activity, as well as reduced H_2_O_2_ release compared to AL controls. Inhibition of mitochondrial Na^+^/Ca^2+^ exchange abolished the mitochondrial adaptive effects of CR, eliminating its protection against damage in cardiomyocytes and perfused hearts. These findings demonstrate that CR protects the heart from I/R injury by enhancing CGP-sensitive Na^+^-dependent mitochondrial Ca^2+^ efflux, preserving mitochondrial function, limiting redox imbalance, and improving post-ischemic cardiac recovery.

**GRAPHICAL ABSTRACT:** 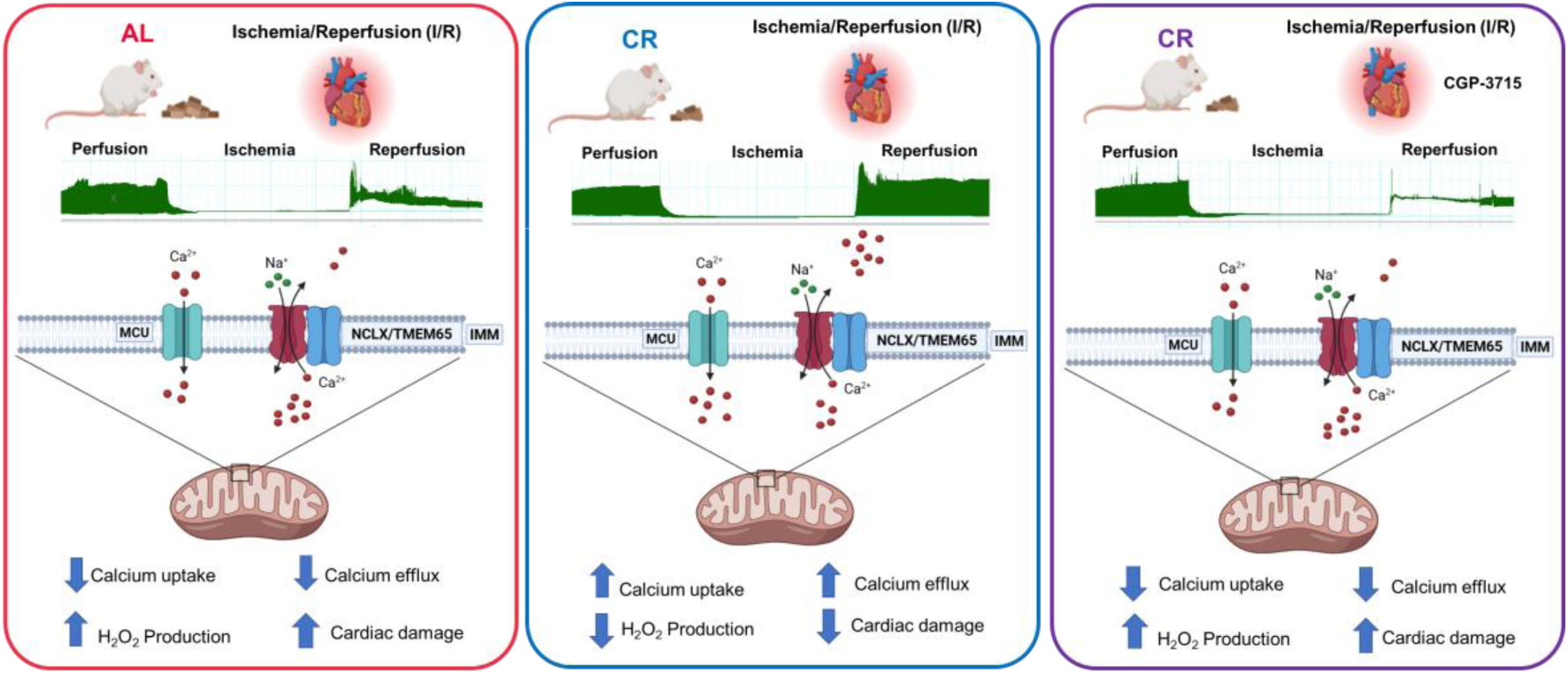

## 1. INTRODUCTION

Ischemic heart disease is a pathological condition characterized by the interruption of blood flow to the heart, and remains one of the leading causes of mortality and morbidity worldwide (Lu et al., 2015; Virani et al., 2020; GBD, 2025; GBD, 2026). The lack of blood supply results in an imbalance between metabolic demands and oxygen delivery, causing tissue hypoxia. Restoration of blood flow, although essential for tissue salvage, paradoxically initiates a series of deleterious events collectively known as ischemia/reperfusion (I/R) injury (Jennings & Reimer, 1981; Del Re et al., 2019; Heusch, 2020). Reperfusion disrupts ion homeostasis, causing cytosolic Ca^2+^ overload, hypercontracture, and mitochondrial Ca^2+^ accumulation (Duchen et al., 1993; Bompotis et al., 2016). These events, together with excessive reactive oxygen species (ROS) generation (Cadenas, 2018) promoted by excessive mitochondrial Ca^2+^, promote opening of the mitochondrial permeability transition pore (mPTP), due to mitochondrial inner membrane protein thiol oxidation (Vercesi et al., 2018). This non-selective inner membrane permeabilization leads to dissipation of the inner membrane potential and loss of oxidative phosphorylation, as well as the release of apoptogenic factors (Hurst et al., 2017; Vercesi et al., 2018; Garbincius and Elrod, 2022). Reoxygenation also promotes redox modifications in proteins, lipids, and DNA, inducing ferroptotic and necrotic cell death (Cao et al., 2006; Eltzschig and Eckle, 2011; Heusch, 2020).

The importance of mitochondrial Ca^2+^ management in cardiac injury is well established, as modulation of mitochondrial calcium transport profoundly affects cardiac outcomes (Garbincius and Elrod, 2022). Mitochondrial Ca^2+^ uptake is mediated by the mitochondrial calcium uniporter MCU (Baughman et al., 2011; De Stefani et al., 2011; Feno et al., 2021), while Ca^2+^ efflux in heart occurs predominantly through the mitochondrial Na⁺/Ca^2+^ exchanger (mNCX), whose molecular identity has recently become the focus of debate. Although the NCLX protein has been widely considered the main mediator of mitochondrial Ca^2+^ extrusion, new data suggest TMEM65 contributes to this process and could represent an essential component of the exchanger machinery (Palty et al., 2010; Palty et al., 2012; Luongo et al., 2017; Garbincius et al., 2022; Vetralla et al., 2025; Zhang et al., 2025; Zhang et al., 2026a; Zhang et al., 2026b; see a thoughtful discussion by Garbincius and Elrod, 2025).

Genetic studies emphasize the importance of mitochondrial Ca^2+^ transport in myocardial infarction, demonstrating that MCU deletion can attenuate I/R injury (Pan et al., 2013). Conversely, loss of NCLX causes rapid cardiac dysfunction and death, while enhanced mitochondrial Ca^2+^ extrusion protects against ischemic injury (Luongo et al., 2017; Garbincius et al., 2022), indicating that limiting mitochondrial Ca^2+^ overload protects the ischemic heart.

Caloric restriction (CR) is also known to protect against cardiac injury induced by I/R, and previous studies indicate it acts by modulating mitochondrial function, improving redox balance, and limiting the production of oxidants (Lanza et al., 2012; David et al., 2018; Guo et al., 2023). During CR, antioxidant defence systems are activated, decreasing intracellular oxidant levels across multiple tissues (Cerqueira et al., 2011; Lanza et al., 2012). In addition, we found that CR protects against Ca^2+^-induced mPTP opening in brain and liver (Amigo et al., 2017; Menezes-Filho et al., 2017), although this effect was not initially observed in the heart (Serna et al., 2020). This apparent tissue-specific discrepancy may reflect differences in experimental conditions, as mPTP induction was only verified using mitochondria isolated under control conditions, and changes could become evident specifically under pathological states such as I/R. Alternatively, protection against Ca^2+^ overload in heart may depend less on mitochondrial Ca^2+^ uptake and more on efflux mechanisms, which are inactive in the sodium-free media commonly used in studies with isolated mitochondria.

Based on these observations, we hypothesized that CR may confer protection against cardiac I/R injury by enhancing mitochondrial Ca^2+^ extrusion, thereby limiting mitochondrial Ca^2+^ overload and preventing mPTP opening, possibly by enhancing mNCX activity. To test this hypothesis, we quantitatively examined the effects of CR on mitochondrial Ca^2+^ transport activity and I/R injury using complementary animal and cell models of CR.

## 2. MATERIALS AND METHODS

### 2.1. Rat Caloric Restriction and Preparation of Sera

All procedures were approved by the local ethics committee (*Comissão de Ética em Uso Animal*, CEUA IQ-USP, protocol #309) under Brazilian bylaws in general accordance with the NIH Guide for the Care and Use of Laboratory Animals. To minimize total animal usage in the absence of known sex-specific effects, only male Sprague-Dawley rats were used, complying with ethical recommendations regarding animal reduction (while US funding bodies and oversight committees often mandate the use of both male and female animals, local frameworks prioritize minimizing total animal counts when sex-specific differences are not central to the study). Eight-week-old rats were randomly divided into two groups: *ad libitum*-fed (AL) rats receiving the AIN-93G diet and caloric restriction (CR) rats receiving 40% less AIN-93G diet supplemented with micronutrients to maintain equivalent vitamin and mineral intake (Cerqueira and Kowaltowski, 2010), resulting in lower weight gain (**Figure S1**). Dietary intake in AL animals was quantified weekly, and CR amounts adjusted accordingly. Animals had free access to water throughout the study. Rats were euthanized with a lethal intraperitoneal dose of ketamine (100 mg/kg), xylazine (10 mg/kg), and acepromazine (6 mg/kg) after 16 weeks of dietary intervention. Blood and hearts were collected, and serum was obtained by centrifugation at 300 x g for 20 min and stored at -80°C. Immediately before use, serum aliquots were thawed, pooled with five animals per group, and heat-inactivated at 56°C for 30 min.

### 2.2. Cardiac Ischemia/Reperfusion and Functional Assessment

After 16 weeks of the dietary intervention, animals anesthetized and euthanized with intraperitoneal ketamine (100 mg/kg), xylazine (10 mg/kg), and acepromazine (6 mg/kg) plus sodium heparin (1000 units/kg) had the aorta rapidly cannulated for perfusion using a Langendorff heart perfusion system. Hearts were removed and the aorta cannulated in under 3 min. Hearts were retrograde-perfused using a Langendorff apparatus (AD Instruments) with oxygenated gas mixture bubbled with 95% O_2_. All perfusions were at a constant flow of 12 mL. min^−1^ at 37°C. Hearts were perfused for 20 min before undergoing 40 min of global ischemia; reperfusion was performed for 30 min. The perfusion solution consisted of Krebs buffer containing (in mM): 118 NaCl, 25 NaHCO_3_, 1.2 KH₂PO₄, 4.7 KCl, 1.2 MgSO₄, 1.25 CaCl₂, 10 glucose, and 10 HEPES, pH 7.4, at 37°C, supplemented with either 0.5 μM CGP-37157 (CGP) to inhibit mNCX or an equal volume (0.1%) of the solvent DMSO (control). A water-filled balloon connected to a pressure transducer was inserted into the left ventricle through the left atrium to monitor cardiac hemodynamic function using the PowerLab system (AD Instruments). The balloon volume was adjusted to set an initial left ventricular end-diastolic pressure (LVEDP) below 10 mmHg. The following hemodynamic parameters were evaluated: left ventricular systolic pressure (LVSP), left ventricular end-diastolic pressure (LVEDP), left ventricular developed pressure (LVDP = LVSP − LVEDP), maximal rate of pressure development (dP/dtmax), maximal rate of pressure decline (dP/dtmin), and rate-pressure product (RPP = LVDP × heart rate). At the end of the protocol, hearts were collected for further analyses such as mitochondrial isolation.

### 2.3. Mitochondrial Isolation and Sample Preparation

Cardiac tissue was minced into small pieces, washed twice with ice-cold PBS, and homogenized using a pre-chilled glass Potter homogenizer (20 strokes) in isolation buffer containing 300 mM sucrose, 10 mM K⁺-HEPES (pH 7.2), 1 mM K⁺-EGTA, and 1 mg/mL BSA. Nuclei and cell debris were removed by centrifugation at 800 x g for 3 min. The supernatant was then centrifuged at 9,300 x g for 10 min to pellet mitochondria. The mitochondrial pellet was resuspended in isolation buffer and used immediately (David et al., 2018).

### 2.4. HL-1 Cardiomyocyte Cell Cultures

HL-1 cardiomyocytes were maintained in T-75 flasks at 37°C in a humidified atmosphere containing 5% CO₂. Cells were cultured in Claycomb medium (JRH Biosciences) supplemented with 0.1 mM norepinephrine (Sigma-Aldrich), 100 U/mL penicillin, 100 μg/mL streptomycin, 2 mM glutamine, and 10% fetal bovine serum. Experiments were initiated when cells reached 100% confluence. Intact cell experiments were performed in standard buffer (pH 7.4) containing (in mmol/L): 137 NaCl, 5 HEPES, 22 glucose, 20 taurine, 5 creatine, 5.4 KCl, 1 MgCl₂, 5 pyruvate, and 1 CaCl₂. Permeabilized cell experiments were conducted in the same experimental buffer used for isolated mitochondria, described below.

### 2.5. Mitochondrial Ca^2+^ Uptake Measurements

Mitochondrial Ca^2+^ uptake was assessed in digitonin-permeabilized cells (1 . 10^6^ dispersed cells in the presence of 3 μM digitonin) or isolated cardiac mitochondria (250 μg/mL) using 0.1 μM Calcium Green-5N (Molecular Probes). Fluorescence was monitored in a Hitachi F4500 spectrofluorometer (Ex = 506 nm, Em = 531 nm, slit width = 5 nm) at 37°C, in 2 mL experimental media containing 125 mM sucrose, 65 mM KCl, 10 mM HEPES, 2 mM K^+^ phosphate, 2 mM MgCl_2_, 200 μM ATP, and 0.2% BSA; pH was adjusted to 7.2 with KOH, plus 1 mM malate and 1 mM glutamate as substrates. Additionally, 50 µM Ca^2+^, 0.5 µM RuR, 5 µM CGP and 20 mM NaCl were added as indicated in the figures. Since Calcium Green-5N binds extramitochondrial Ca^2+^ reversibly, mitochondrial calcium uptake results in decreased fluorescence intensity (Amigo et al., 2017; Serna et al., 2022a), providing quantitative measurements that are not limited by intramitochondrial precipitation. Sequential Ca^2+^ pulses were added until mitochondria no longer exhibited evidence of calcium uptake. Total calcium retention capacity was calculated as the sum of all calcium additions, with fluorescence changes calibrated with known calcium additions.

### 2.6. Mitochondrial Ca^2+^ Extrusion Activity

Na^+^-dependent and -independent mitochondrial Ca^2+^ release activity was evaluated in digitonin-permeabilized cells and isolated cardiac mitochondria using Calcium Green-5N fluorescence, following protocols previously established by our group (Serna et al., 2022b). Briefly, mitochondria or cells were incubated as for Ca^2+^ uptake, above, and loaded with one pulse of 50 µM calcium in experimental media identical to that used in uptake measurements. RuR (0.5 μM) was then added to promote inhibition of uptake, in the presence of 5 µM CGP and 20 mM NaCl, where indicated. Extrusion rates were calculated from the linear fluorescence increases in the first 200 seconds following RuR addition.

### 2.7. Mitochondrial H_2_O_2_ Release Detection

H_2_O_2_ release from isolated mitochondria in experimental media under the same conditions as Ca^2+^ uptake measurements, using the Amplex Red assay. Amplex Red (1 μM; Molecular Probes), in the presence of horseradish peroxidase (HRP, 1 U/mL), is converted into the fluorescent product resorufin in a 1:1 stoichiometric reaction with H_2_O_2_. Fluorescence was monitored over time using a Hitachi F4500 spectrofluorometer at 37°C with continuous stirring (Ex = 563 nm, Em = 587 nm) for data acquisition, during 10 min. Data were calibrated with known additions of commercial H_2_O_2_.

### 2.8. Oxygen Consumption Measurements

Oxygen consumption was measured using a High Resolution Oroboros Oxygen electrode system under the same conditions as calcium uptake, in the experimental buffer described above, with 20 mM NaCl, 50 μM CaCl₂, 1.25 mM EGTA, 0.5 μM RuR, and 5 μM CGP added as indicated. Data shown in the main manuscript were conducted under State 4 conditions, in the presence of 1 μM oligomycin. ADP (1 mM, State 3) and 1.25 μM CCCP (State 3u) were added to induce oxidative phosphorylation and maximize electron transport in supplementary figures.

### 2.9. Caloric Restriction versus *Ad Libitum* Model in Cardiac Cells

After establishment of cell cultures, Claycomb medium supplemented with fetal bovine serum was replaced with media containing 10% heat-inactivated rat serum obtained from AL or CR animals, prepared as described above (Cerqueira et al., 2012; Cerqueira et al., 2016; de Cabo et al., 2003). Cells were incubated for 24 h before experiments were performed.

### 2.10. Simulated Ischemia/Reperfusion in Cardiomyocytes

Simulated ischemia/reperfusion was induced in cardiac HL-1 cells (∼1.5 × 10⁵ cells/mL) by metabolic inhibition using 500 μM KCN and 2 mM 2-deoxyglucose in glucose- and pyruvate-free standard buffer for 50 min, followed by removal of KCN and re-introduction of substrates, as previously described (Facundo et al., 2006; Facundo et al., 2007). Cells were pretreated with experimental drugs for 24 h before simulated ischemia. Simulated reperfusion was induced by replacing the ischemic medium with control buffer containing glucose and pyruvate for 90 min, imitating many of the mitochondrial effects of I/R seen in heart (Facundo et al., 2006; Facundo et al., 2007). Cell death was measured using 7.5 µM propidium iodide staining.

### 2.11. Western Blots

Cells or isolated mitochondria were lysed over ice in the presence of RIPA buffer containing protease and phosphatase inhibitor cocktails, and total protein levels were quantified using the Pierce BCA Protein Assay kit. Lysates were prepared in sample buffer (2% SDS, 10% glycerol, 0.062 M Tris pH 6.8, 0.002% bromophenol blue, 5% 2-mercaptoethanol), 40 μg of total protein was loaded onto 10% SDS/PAGE, and electro-transferred to PVDF membranes with 0.2 µm. Membranes were blocked with 5% BSA in TTBS (20 mM Tris pH 7.5, 150 mM NaCl, and 0.1% Tween 20) for 1 h at room temperature before incubation with Total OXPHOS Human WB Antibody Cocktail (Abcam, Cambridge, UK:110411, 1:1000), MCU (Cell Signaling: 14997, 1:1000), MCUB (Sigma: HPA048776, 1:500), MICU1 (Sigma: HPA037480, 1:1000), MICU2 (Sigma: HPA045511, 1:500), MICU3 (Sigma: HPA024771, 1:250), EMRE/SMDT1 (Sigma: HPA060340, 1:500), NCLX/SLC24A6 (Proteintech: 21430-1AP, 1:1000), or TMEM65 (Abcam: Ab236861, 1:1000) antibodies; primary antibodies were incubated at 4°C overnight. Fluorescent secondary anti-mouse or anti-rabbit antibodies (1:10 000) were incubated for 1 h at room temperature prior to 660 nm fluorescence detection using Odyssey ® CLx (Li-Cor). Quantification of band densitometry was performed using the ImageLab software. Since no single protein is a reliable housekeeping control in metabolic contexts, protein expression values obtained were normalized per pixel of total protein in each well, as evidenced by Ponceau S dye (Sigma, See Supplementary material for uncropped images of normalization stains and of the blots quantified).

### 2.12. Data Analysis

Results are presented as means ± SD unless indicated in the legend, with individual symbols depicting data from biological replicates (defined as data obtained from distinct animal individuals or distinct cell preparations from a distinct experimental day). Technical replicates, when present, were analyzed when pooled, as part of data within each single biological replicate. Statistical analyses were performed using GraphPad Prism^®^ 8 t test (for comparison between 2 samples) and ANOVA multiple comparisons, while representative graphs were generated using Origin 2025. Langendorff analysis was conducted using LabChart Reader software. Calcium efflux analyses were performed using the Mitochondrial Calcium Efflux Calculator software generated in-house for this purpose (https://github.com/hebertyfacundo/NCLXapp) and Microsoft Excel. Outliers were identified using Grubbs’ test (α = 0.05) only for oxygen consumption measurements (data without outlier removal also did not show statistical differences). Statistical significance was considered at p ≤ 0.05.

## 3. RESULTS

### 3.1. Caloric Restriction Prevents Cardiac Dysfunction Following Ischemia/Reperfusion

To determine whether weight gain limitation promoted by caloric restriction (CR, **Figure S1**) protects against ischemia/reperfusion (I/R)-induced cardiac injury, hearts from *ad libitum*-fed (AL, in red) and CR (in blue) animals were Langendorff-perfused and subjected to *ex vivo* global I/R (schematic in **Figure 1A**). Reperfusion markedly increased left ventricular end-diastolic pressure (LVEDP) in AL hearts, whereas CR hearts maintained LVEDP values close to baseline, indicating substantial preservation of diastolic function (**Figure 1B**), and thus marked cardioprotection. Left ventricular systolic pressure (LVSP) did not differ significantly between groups (**Figure 1C**). Left ventricular developed pressure (LVDP) was preserved in CR hearts after I/R, but markedly decreased in AL hearts after reperfusion (**Figure 1D**). Similarly, the rate-pressure product (RPP), an indicator of cardiac mechanical performance, remained near preischemic baseline in CR hearts, but declined significantly in AL hearts (**Figure 1E**). Together, these findings demonstrate that CR markedly preserves cardiac function following I/R.

**Figure 1.**
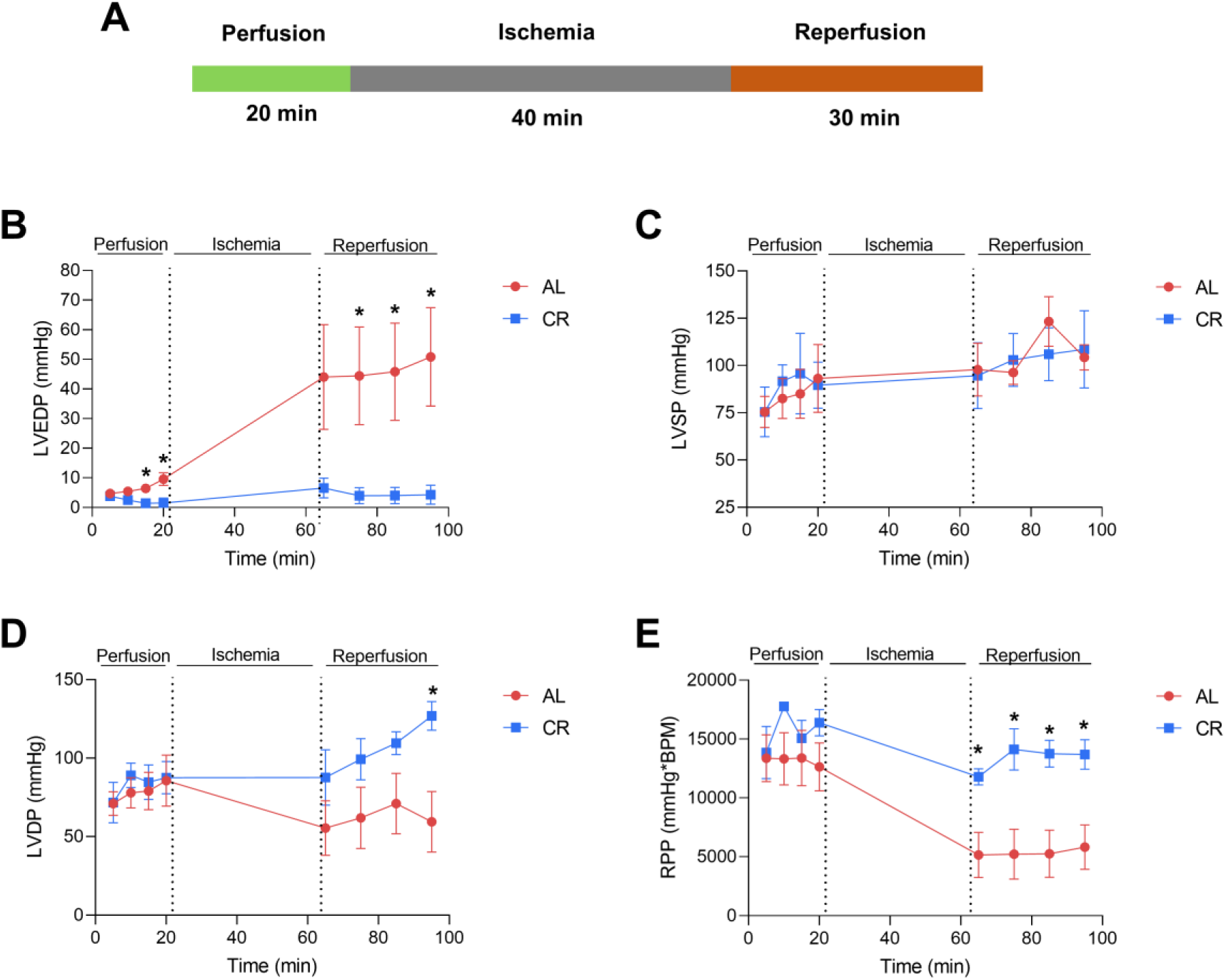
Caloric restriction prevents cardiac damage after ischemia and reperfusion. (A) Schematic of the heart perfusion protocol. (B) LVEDP (Left Ventricular End-Diastolic Pressure). (C) LVSP (Left Ventricular Systolic Pressure). (D) LVDP (Left Ventricular Developed Pressure). (E) RPP (Rate Pressure Product). Statistical significances were determined by one-way ANOVA. Data are expressed as averages ± SE of 4 different individuals in each experimental group. * = p < 0.05.

### 3.2. CR Increases Mitochondrial Ca^2+^ Retention Capacity and Decreases H_2_O_2_ Release

Mitochondrial permeability transition pore (mPTP) opening, limiting mitochondrial Ca^2+^ retention capacity, is a major contributor to cardiac injury during I/R (Duchen et al., 1993; Griffiths and Halestrap, 1995; Di Lisa et al., 2003; Patel et al., 2024). To investigate mitochondrial mechanisms underlying CR-induced cardioprotection, cardiac mitochondria were isolated from AL and CR animals after 16 weeks of dietary intervention, and their ability to accumulate and retain Ca^2+^ was assessed by monitoring extramitochondrial Ca^2+^ with the membrane-impermeable fluorescent probe Calcium Green (**Figure 2A**), a technique that, unlike loading with intramitochondrial probes, allows for total mitochondrial calcium accumulation quantification, as it is not limited by matrix ion precipitation. During the assay, isolated mitochondria were challenged with sequential pulses of 50 μM Ca^2+^ every 200 seconds. Each Ca^2+^ addition produced a transient increase in fluorescence, reflecting elevated extramitochondrial Ca^2+^, followed by a progressive decline as mitochondria accumulated the ion. This process continued until the mitochondrial calcium retention capacity (CRC) was exceeded, triggering mPTP opening and the abrupt release of accumulated Ca^2+^ back into the medium, detected as an increase in Calcium Green fluorescence at the end of the time scan (Chinopoulos et al., 2003; Serna et al., 2020).

**Figure 2.**
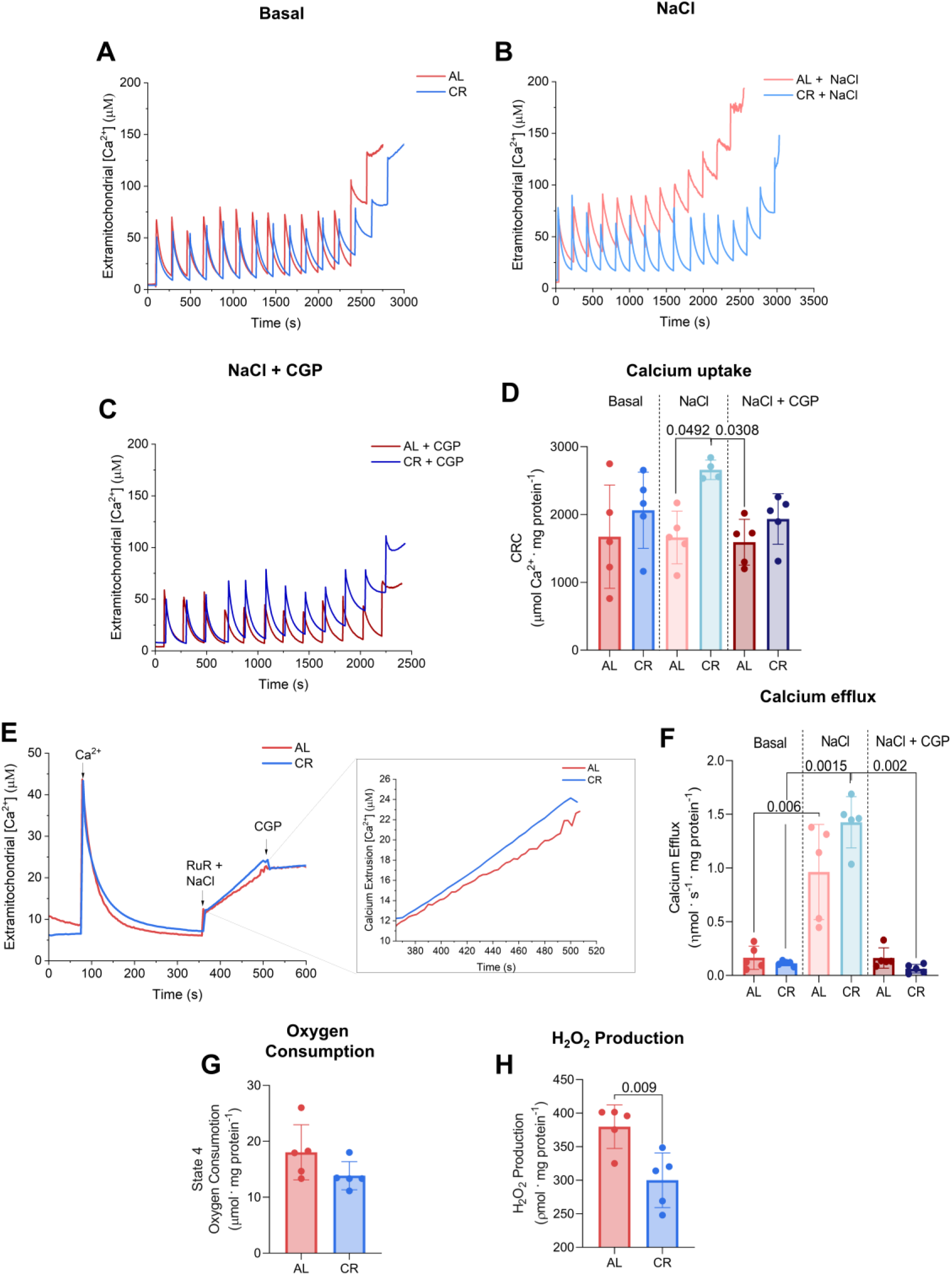
Caloric restriction increases Na^+^-sensitive Ca^2+^ retention capacity and decreases H_2_O_2_ release. (A-C) Representative graphs of calcium uptake traces using malate plus glutamate as substrates and Calcium Green-5N as an extramitochondrial probe, as described in the Methods. Ca^2+^ boluses (50 µM) were added every 200 seconds, generating the upward fluorescence peaks, followed by mitochondrial uptake, decreasing fluorescence until retention capacity was reached. (A) Basal conditions, (B) presence of 20 mM NaCl, (C) 20 mM NaCl plus 5 μM CGP. (D) Calcium uptake quantifications from traces such as those in panels A-C. (E) Representative trace of calcium efflux assessment as described in Methods; 50 µM Ca^2+^, 0.5 μM RuR, 5 µM CGP and 20 mM NaCl were added as indicated. The insert is a magnified region where data were quantified. (F) Calcium efflux quantification. (G) Basal oxygen consumption, measured as described in Methods. (H) Basal mitochondrial H_2_O_2_ release, measured as described in Methods. Statistical significance was determined by one-way ANOVA. Data are expressed as averages ± SD of 5 different experiments, and individual symbols represent biological replicates. P values are indicated in the graph when < 0.05.

Interestingly, CR and AL mitochondria displayed similar Ca^2+^ uptake profiles when measurements were performed in the absence of Na⁺ (**Figure 2A**, quantified in **Figure 2D**), in keeping with prior data (Serna et al., 2020). In contrast, in the presence of Na⁺, which stimulates mitochondrial Na⁺/Ca^2+^ exchange activity (mNCX), CRC was consistently lower in AL compared to CR mitochondria (**Figure 2B,D**). Inhibition of mNCX with CGP-37157 (CGP; **Figure 2C,D**) abolished this difference, reduxing CRC in CR mitochondria to levels comparable to those of AL mitochondria. These findings suggest that the enhanced CRC and prevention of mPTP observed in CR mitochondria results from increased Na⁺/Ca^2+^ exchange (mNCX) activity.

To directly measure mNCX activity, a Ca^2+^ extrusion protocol was designed (**Figure 2E**; Serna et al., 2022b). Mitochondria were first loaded with a single Ca^2+^ pulse and allowed to complete Ca^2+^ uptake. Efflux was then initiated by adding the MCU inhibitor ruthenium red (RuR) to prevent further Ca^2+^ uptake, in the presence or absence of Na⁺ ions or the mNCX inhibitor CGP. Ca^2+^ efflux rates were determined from the increase in extramitochondrial fluorescence (**Figure 2E**, inset) and quantified in **Figure 2F**. Ca^2+^ extrusion by cardiac mitochondria was largely Na⁺-dependent, as expected (Luongo et al., 2017), and showed a trend toward higher efflux rates in CR mitochondria compared with AL. Changes in mitochondrial Ca²⁺ transport observed between AL and CR animals were not attributable to differences in oxidative phosphorylation, as oxygen consumption (**Figure 2G**) under the conditions measured and other respiratory parameters (**Figure S2**) were indistinguishable between groups. In contrast, steady-state H_2_O_2_ production, a key promoter of protein thiol oxidation and mPTP opening (Vercesi et al., 2018), was significantly lower in CR compared to AL mitochondria (**Figure 2H**), in keeping with prior studies (Colom et al., 2007). Together, these findings indicate that, without altering respiratory function, CR selectively remodels rat heart mitochondrial Ca^2+^ handling and redox status, creating resilience toward mPTP opening.

### 3.3. Serum From CR Animals Increases Ca^2+^ Retention and Efflux in Cardiomyocytes

In prior studies (Cerqueira et al., 2016; Munhoz et al., 2023), we found that many of the metabolic adaptations observed in tissues from CR animals were reproduced in cultured cells exposed to sera collected from these animals, indicating that circulating factors induced by CR are sufficient to acutely confer much of the phenotype. This experimental approach reduces animal use and also minimizes the biological variability inherent to individual animals, while allowing mechanistic studies under controlled conditions. To determine whether enhanced mitochondrial Ca^2+^ transport is similarly transferable, HL-1 cardiomyocytes were incubated for 24 h with serum from either AL or CR rats prior to assessment of mitochondrial Ca^2+^ handling in digitonin-permeabilized cells (see Methods; Serna et al., 2022b; **Figure 3A**).

**Figure 3.**
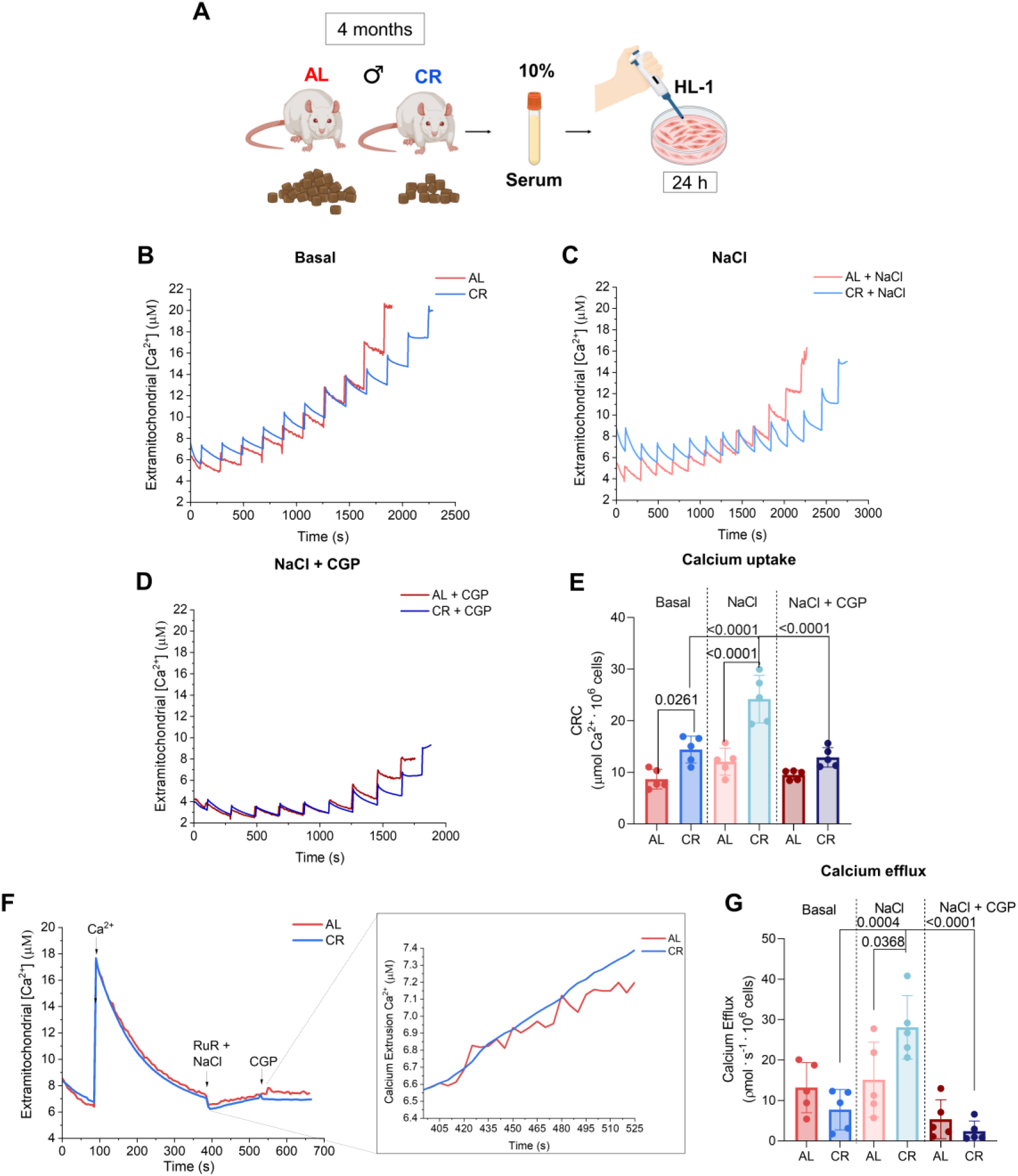
Cells treated with serum from CR animals display increased mitochondrial calcium retention and efflux. (A) Experimental schematic. (B-D) Representative graphs of basal, NaCl, NaCl + CGP calcium uptake in permeabilized cells (see Methods) under conditions similar to Figure 2. (E) Calcium uptake quantification. (F) Representative calcium efflux trace; the insert is a magnification. (G) Calcium efflux quantification. Statistical significance was determined by one-way ANOVA. Data are expressed as averages ± SD of 5 different experiments, and individual symbols represent biological replicates. P values under 0.05 are indicated in the graph.

Consistent with results obtained in hearts, mitochondria from cardiomyocytes in CR serum displayed a greater CRC than those exposed to AL serum under basal conditions (**Figure 3B-D**). This difference was most pronounced in the presence of added Na⁺ (it should be noted that traces of contaminating sodium may be present in permeabilized cell preparations even in the absence of added ion). Inhibition of mNCX with CGP abolished the increase in CRC induced by CR serum (**Figure 3D,E**), indicating that enhanced CGP-sensitive Na⁺-dependent Ca^2+^ extrusion underlies improved CRC and mPTP prevention in CR sera-incubated cells.

To directly assess mNCX activity, mitochondrial Ca^2+^ efflux was measured as described for Figure 2. Cells treated with CR serum exhibited faster Ca^2+^ extrusion than AL-treated cells (**Figure 3F,G**) only in the presence of added Na^+^. CGP eliminated this difference, confirming that CR serum enhances mNCX-mediated Ca^2+^ efflux in cardiomyocyte mitochondria. No differences were observed in oxygen consumption rates measured under different conditions using extracellular flux analysis, with or without pharmacological modulation of MCU or mNCX (**Figure S3**). Together, these findings demonstrate that circulating factors induced by CR are sufficient to recapitulate enhanced mitochondrial Ca^2+^ extrusion and retention observed *in vivo*, supporting a central role for mNCX activation in CR-induced cardioprotective mitochondrial remodelling.

### 3.4. CR Increases NCLX and TMEM65 Protein Expression

To investigate whether alterations in the amounts of mitochondrial calcium transport proteins could explain the functional changes observed, proteins involved in oxidative phosphorylation, mitochondrial calcium uptake and extrusion were quantified in cells and heart mitochondria. No significant differences were detected in electron transport chain proteins in either model (**Figure S4**), consistent with the absence of changes in mitochondrial oxygen consumption rates. Likewise, no differences were observed in the expression of MCU complex components (MCU, MICU1, MICU2, MICU3, MCUB, and EMRE) in either isolated mitochondria (**Figure 4A-I**) or cardiomyocytes (**Figure 4J-Q**; EMRE was undetectable in the cardiomyocytes). Conversely, in isolated cardiac mitochondria, mNCX component TMEM65 was significantly increased, while NCLX (both the 64 KDa and 50 KDa bands) showed a strong trend toward increased expression (p = 0.05; **Figures 4H,I**). In cardiomyocytes, both 64 kDa NCLX and TMEM65 protein expression were increased (**Figure 4P,Q**). This demonstrates that CR and CR serum treatment specifically enhance protein quantities of mNCX components.

**Figure 4.**
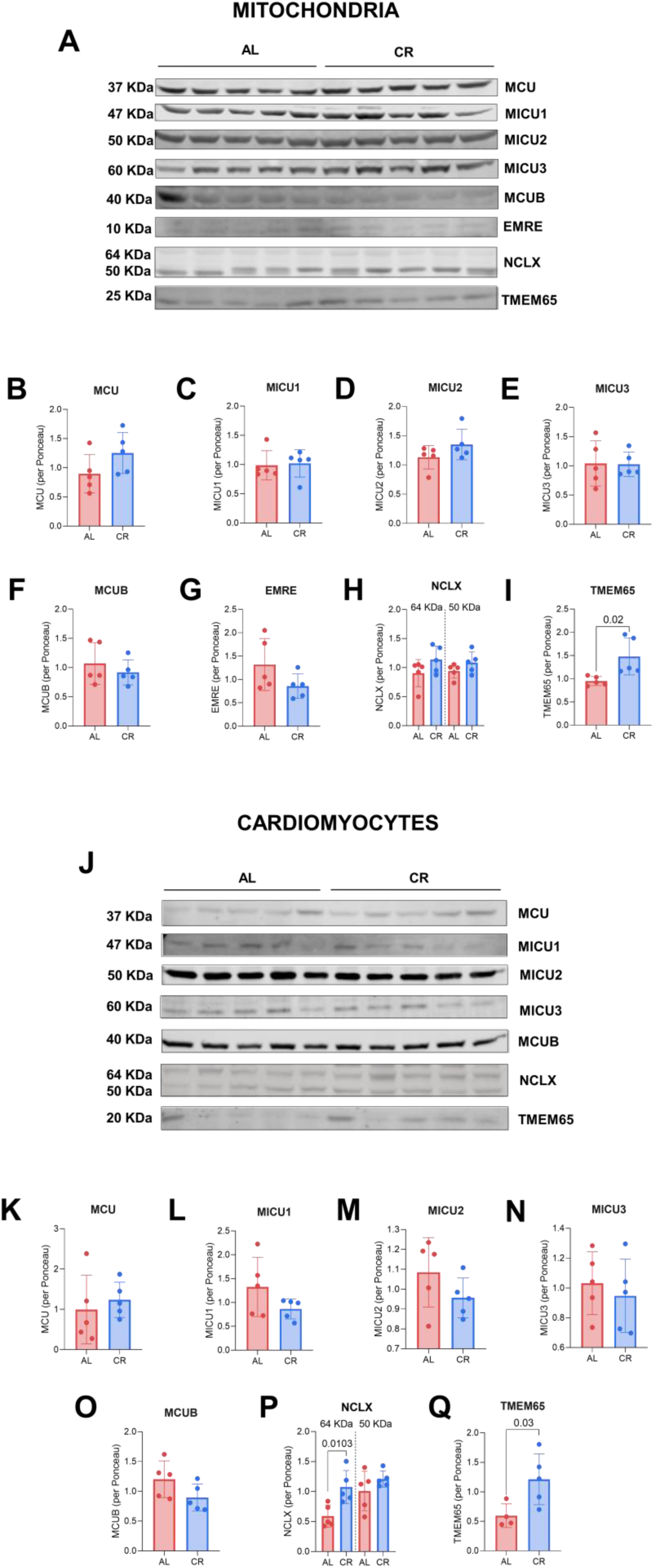
CR increases the amount of NCLX and TMEM65 proteins in hearts and cardiomyocytes. (A and J) Representative western blot images of mitochondrial calcium transport proteins (MCU, MICU1, MICU2, MICU3, MCUB, EMRE, NCLX, and TMEM65) in mitochondria and cells. (B-I and K-Q) Protein quantification (MCU, MICU1, MICU2, MICU3, MCUB, EMRE, NCLX and TMEM65) normalized per Ponceau (see Supplementary Materials for the membrane staining/loading control images). Statistical significance was determined by one-way ANOVA. Data are expressed as averages ± SD of 5 different experiments, and individual symbols represent biological replicates. P values < 0.05 are indicated in the graph.

### 3.5. Inhibition of mNCX Abolishes the Protective Effects of CR in Cardiomyocytes

Given the marked increase in mNCX activity and the expression of its associated proteins (NCLX and TMEM65) in cardiomyocytes exposed to CR serum, together with the established role of mNCX in determining susceptibility to I/R injury (Luongo et al., 2017; Garbincius et al., 2022), we investigated if enhanced mitochondrial Ca^2+^ extrusion underlies CR-mediated cardioprotection. To do so, simulated I/R by metabolic inhibition (see Methods) was induced in cardiomyocytes pretreated with serum from AL or CR animals (**Figure 5**). Cells exposed to CR serum exhibited a strong reduction in cell death compared with AL serum-treated controls (**Figure 5A,B**), confirming that circulating factors induced by CR are sufficient to confer protection. Importantly, pharmacological inhibition of Ca^2+^ extrusion with CGP completely abolished this protective effect, increasing cell death to levels comparable to those observed in AL-treated cells. These findings demonstrate that enhanced CGP-sensitive mitochondrial Ca^2+^ extrusion pathway is required for the cardioprotective effect conferred by CR serum, identifying increased mNCX activity as a key mechanistic mediator of CR-induced resistance to simulated I/R injury in these cells.

**Figure 5.**
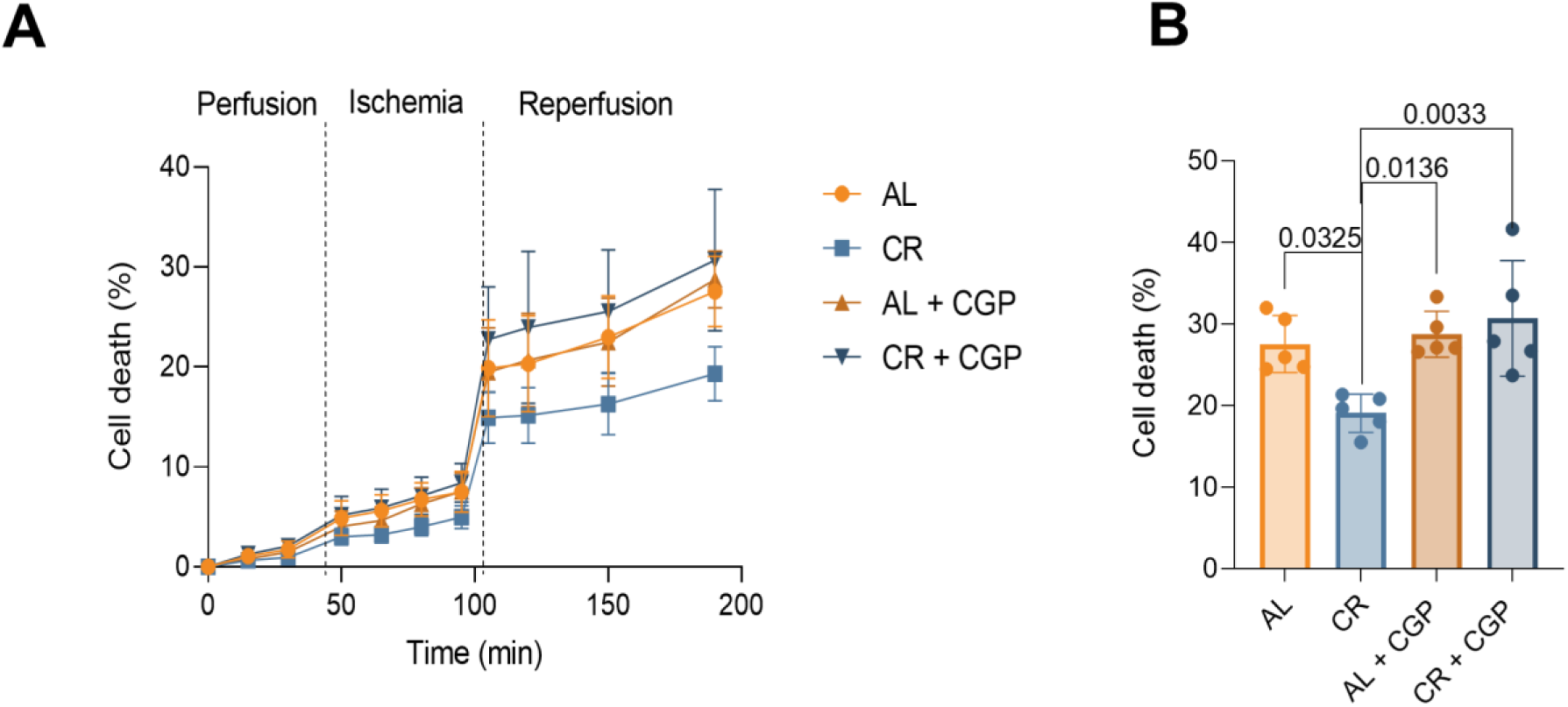
Serum from calorie-restricted animals prevents cell death induced by cyanide/glycemia-simulated ischemia/reperfusion. (A) Graph representing death promoted by simulated ischemia and reperfusion (see Methods). (B) Quantification of cell death after ischemia and reperfusion. Data represent mean ± SD, and individual symbols represent biological replicates. One-way ANOVA statistical tests were used, and p values < 0.05 are indicated in the graphs.

### 3.6. CR Primes Mitochondrial Na^+^/Ca^2+^ Exchange Activity After Ischemia/Reperfusion in Hearts

The results presented thus far demonstrate that CR serum enhances detected mNCX activity in cardiomyocytes, accompanied by increased expression of both NCLX and TMEM65. Importantly, pharmacological inhibition of mNCX abolished the cardioprotective effects conferred by CR serum in cardiomyocytes, indicating that enhanced mitochondrial Ca^2+^ extrusion is required. Consistent with these findings, cardiac mitochondria isolated from CR animals exhibited greater Na⁺-dependent calcium retention capacity, increased TMEM65 expression, and a trend toward higher mNCX activity. Together, these observations, combined with the established cardioprotective role of mNCX (Luongo et al., 2017; Garbincius et al., 2022), led us to hypothesize that CR primes cardiac mitochondria *in vivo* to extrude matrix Ca^2+^ more efficiently during I/R, thereby limiting mitochondrial calcium overload and reducing injury.

To test this, mitochondria were isolated from AL and CR hearts following I/R in a Langendorff perfusion system (**Figure 6A**). Under these conditions, CR mitochondria exhibited significantly greater CRC than AL mitochondria, particularly in the presence of Na⁺ (**Figure 6B-E**). Additionally, mitochondrial Ca^2+^ efflux rates were almost tripled in the CR group compared to AL controls (**Figure 6F,G**). These differences were not attributable to altered respiratory capacity, as oxygen consumption was unchanged between groups (**Figure 6H**). However, respiratory control ratios (RCR, the ratio of oxygen consumption rates in the presence of oxidative phosphorylation to its absence) were higher in CR mitochondria following extramitochondrial Ca^2+^ chelation with EGTA or stimulation of mNCX activity by Na^+^, whereas pharmacological inhibition of mNCX with CGP markedly impaired oxidative phosphorylation efficiency, equalizing CR and AL samples (**Figure S5**). This result is consistent with enhanced mNCX activity in the CR group, protecting against loss of RCRs. Finally, H_2_O_2_ release was significantly lower in CR mitochondria compared to AL after I/R (**Figure 6I**). This redox advantage was lost when mitochondrial Ca^2+^ transport was experimentally manipulated by inhibiting the MCU with RuR or mNCX with CGP, indicating the protective redox effect of CR was Ca^2+^ transport-dependent. Together, these findings indicate that CR preserves mitochondrial calcium homeostasis after I/R through enhanced mNCX-dependent Ca^2+^ extrusion, thereby limiting redox imbalance and maintaining mitochondrial function.

**Figure 6.**
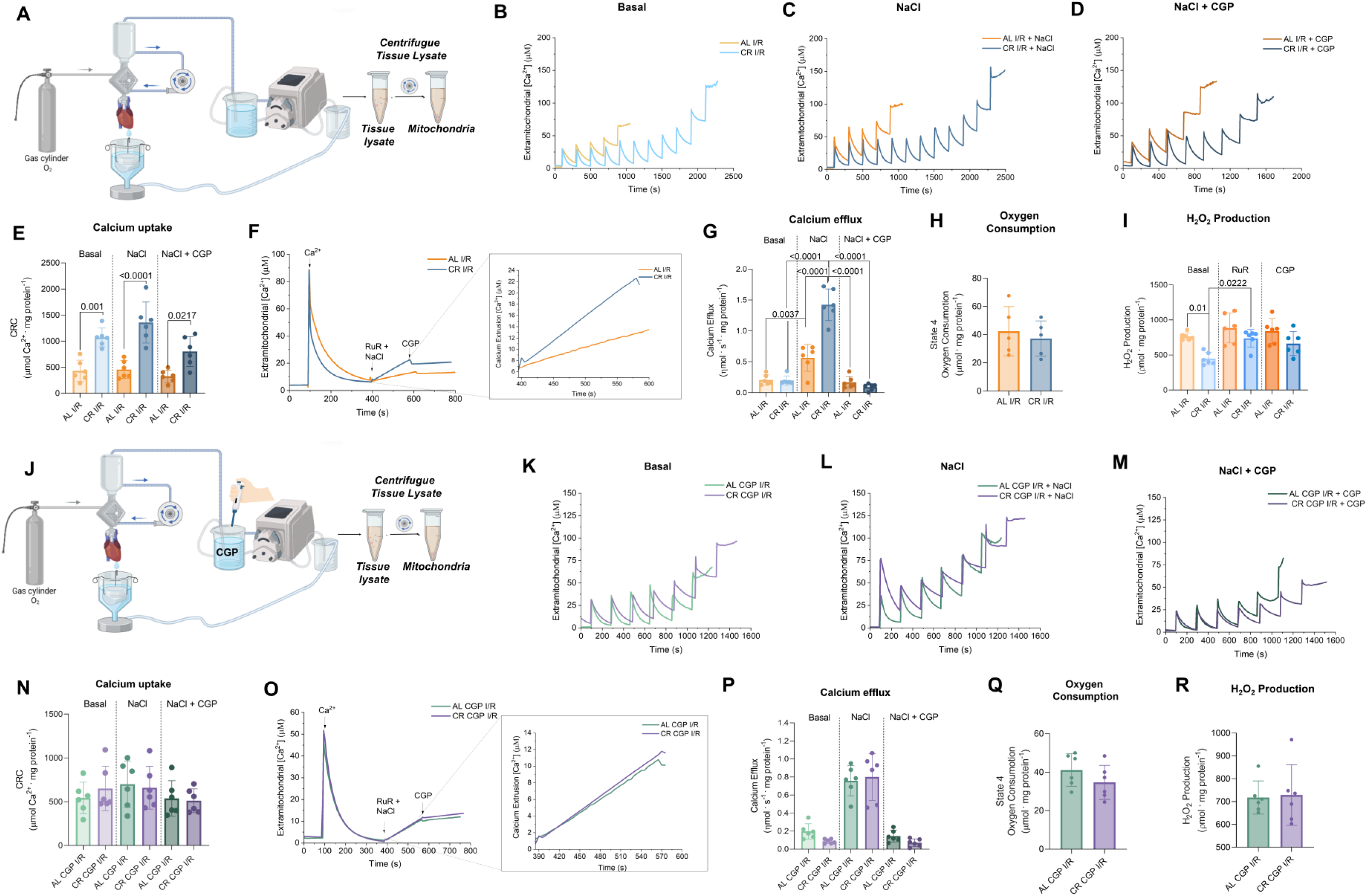
CR increases calcium efflux and decreases H_2_O_2_ production after I/R. (A, J) Experimental schematic of Langendorff I/R with (Panels B-I) or without (Panels K-R) 10 µM CGP in the perfusion buffer. (B-D and K-M) Representative graphs of basal, NaCl, NaCl + CGP calcium uptake using the substrates malate + glutamate, calcium green-5N of mitochondria isolated after I/R (see Methods). (E,N) Calcium uptake quantification. (F,O) Representative graph of calcium efflux. (G,P) Calcium efflux quantification. (H,Q) Basal respiration. (I,R) H_2_O_2_ release under State 4 conditions and in the presence of 0.5 µM RuR and 5 μM CGP, where indicated. Statistical significance was determined by one-way ANOVA. Data are expressed as averages ± SD, and individual symbols represent biological replicates. P values < 0.05 are indicated in the graph.

If the preservation of mitochondrial calcium retention capacity and redox balance following I/R in CR hearts depends on enhanced mitochondrial Na⁺/Ca^2+^ exchange, then inhibition of this pathway during heart perfusion should abolish the protective effects of the diet. To test this, AL and CR hearts were subjected to I/R in the presence of CGP in the heart perfusion buffer (**Figure 6J**). Mitochondria isolated from inhibitor-treated CR hearts lost their enhanced CRC, including under Na⁺-stimulated conditions (**Figure 6K-N**). Likewise, mitochondrial Ca^2+^ efflux was reduced to levels comparable to those observed in AL hearts (**Figure 6O,P**), although Na^+^-induced enhanced extrusion was present, indicating CPG added during the perfusion was effectively washed out by mitochondrial isolation. Basal respiration remained unchanged (**Figure 6Q**), and the improvement in oxidative phosphorylation efficiency (respiratory control ratios) previously observed in CR mitochondria was abolished by CGP treatment during perfusion (**Figure S6E**). Furthermore, inhibition of mNCX during perfusion increased H_2_O_2_ release from CR mitochondria to levels equal to those of AL controls (**Figure 6R**). These findings demonstrate that enhanced mitochondrial Na⁺/Ca^2+^ exchange during I/R is required to preserve mitochondrial calcium homeostasis, oxidative phosphorylation, and redox balance in hearts from CR animals.

### 3.7. Inhibition of mNCX Abolishes the Cardioprotective Effects of CR

Results so far suggest that CR primes heart mitochondria for enhanced Na^+^-dependent Ca^2+^ efflux during I/R, preventing mPTP. We next sought to verify if these mitochondrial alterations were the cause for functional cardioprotection promoted by CR during I/R in hearts. We found that hearts from CR animals perfused with CGP exhibited a marked increase in LVEDP during reperfusion (**Figure 7A,B**), reaching results similar to those seen in AL hearts, and indicating impaired diastolic relaxation and loss of the protection observed in untreated CR hearts. Consistent with this finding, while LVSP remained similar among all groups (**Figure 7C**), both LVDP (**Figure 7D**) and RPP (**Figure 7E**) were significantly reduced by CGP in CR hearts, demonstrating impaired recovery of cardiac contractile performance. Together, these findings demonstrate that inhibition of the mNCX pathway abolishes both mitochondrial and functional benefits of CR, establishing enhanced mitochondrial Ca^2+^ extrusion as a key mediator of CR-induced cardioprotection against I/R injury.

**Figure 7.**
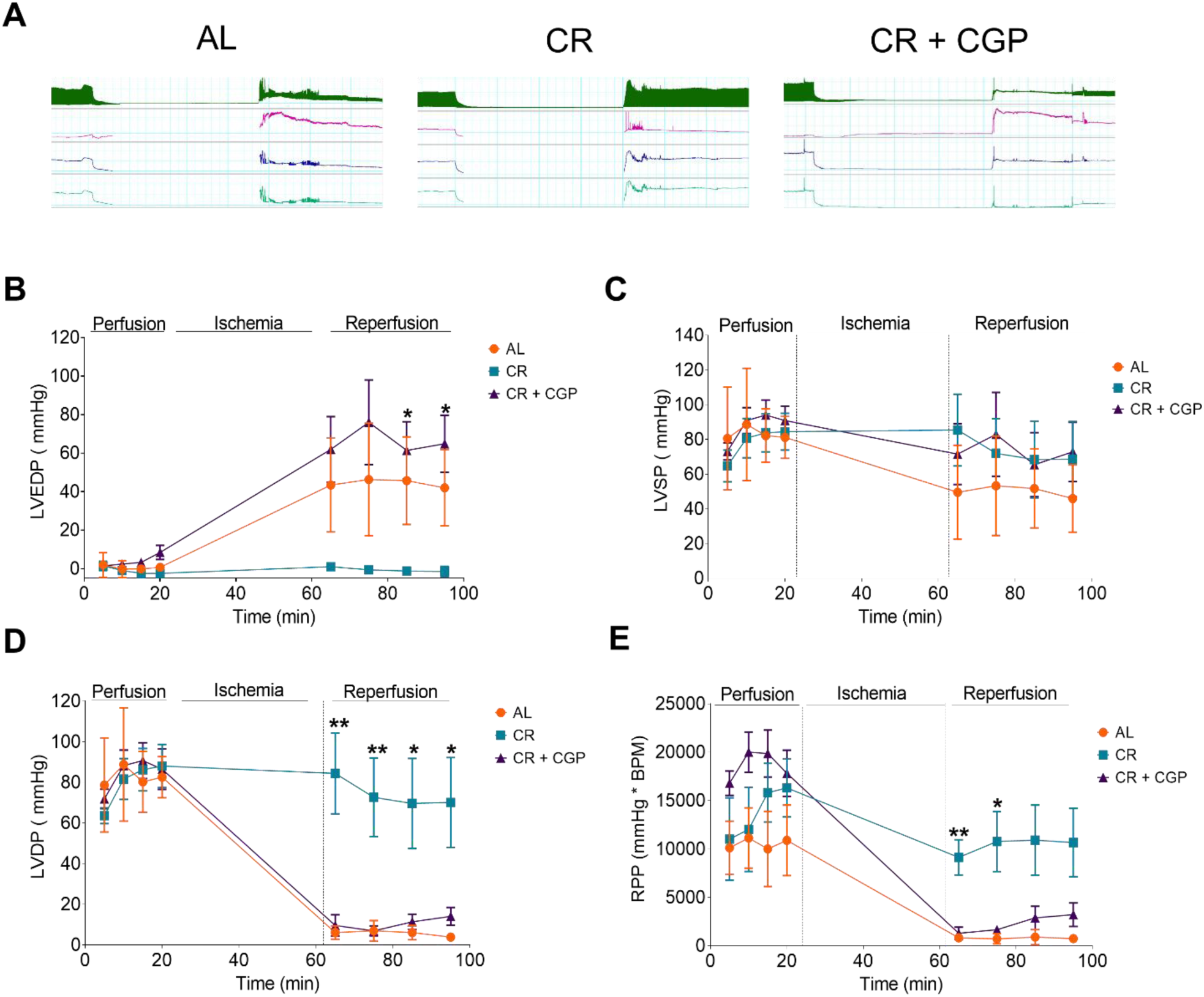
Inhibition of mNCX during I/R in the heart causes CR to lose its protective effect. (A) Representative perfusion, ischemia, and reperfusion traces (AL, CR, CR+CGP), (B) LVEDP (Left Ventricular End-Diastolic Pressure), (C) LVSP (Left Ventricular Systolic Pressure).(D) LVDP (Left Ventricular Developed Pressure). (E) RPP (Rate Pressure Product). Statistical significance was determined by one-way ANOVA. Data are expressed as averages ± SE of 4 different experiments representing distinct individual animals in each group. *P < 0.05; **p < 0.01.

## 4. DISCUSSION

Ischemic heart disease remains one of the leading causes of cardiovascular morbidity and mortality worldwide (GBD, 2025; Virani et al., 2020). Cardiac injury can be markedly attenuated by dietary restriction interventions in rodent models, including intermittent fasting and caloric restriction (Ahmet et al., 2005; Shinmura et al., 2011). Notably, the protective effects of CR extend beyond reducing the incidence of coronary disease risk, as it also increases myocardial resistance to experimental global ischemia/reperfusion injury (Shinmura et al., 2011). Indeed, we find that hearts from rats subjected to 16 weeks of CR are strongly protected against global ischemia (**Figure 1**), exhibiting post-ischemic functional recovery that was nearly indistinguishable from that of non-ischemic hearts. In contrast, hearts from *ad libitum*-fed animals developed marked contractile dysfunction following the same insult. This suggested that CR induces cellular adaptations within the myocardium itself, potentially by increasing mitochondrial resistance to permeability transition pore (mPTP) opening, a Ca^2+^- and redox-sensitive event that is a central determinant of I/R-induced cardiac injury (Guo et al., 2023; Halestrap et al., 2004; Bernardi and Di Lisa, 2015).

Consistent with this hypothesis, CR has been extensively shown to improve mitochondrial redox homeostasis and reduce oxidative stress across multiple tissues (Sohal and Weindruch, 1996; López-Lluch et al., 2006; Madeo et al., 2014). Moreover, our group previously demonstrated that CR modulates mitochondrial Ca^2+^ transport and resistance to mPTP opening in the brain, liver, and kidney (Amigo et al., 2017; Menezes-Filho et al., 2017; Serna et al., 2022). Based on these findings, we hypothesized that CR could protect the heart by modulating mitochondrial Ca^2+^ handling, increasing the threshold for mPTP opening during I/R.

Interestingly, we found that CR markedly increased Ca^2+^ uptake capacity in cardiac mitochondria before mPTP induction, an effect abolished by the mNCX inhibitor CGP (**Figure 2**). Although basal mNCX activity was not significantly altered between groups, mitochondria from CR hearts exhibited increased expression of the mNCX regulatory component TMEM65 (**Figure 4**), suggesting that CR may prime the exchanger for enhanced activity under conditions of Ca^2+^ overload. Indeed, mitochondrial Ca^2+^ retention capacity was identical in CR and AL mitochondria in the absence of Na⁺ or in the presence of CGP (**Figure 2**), demonstrating that the enhanced Ca^2+^ retention induced by CR is dependent on mNCX activity. Additionally, mNCX activity following I/R was markedly higher in CR mitochondria than in AL controls (**Figure 6**), further demonstrating that CR primes the exchanger for enhanced activity under conditions of stress. Also consistent with this finding, the elevated H_2_O_2_ release observed in AL mitochondria after I/R was completely abolished by pharmacological modulation of mitochondrial Ca^2+^ transport (**Figure 6**), indicating that differences in redox state were directly driven by altered mitochondrial Ca^2+^ fluxes. Interestingly, prior results have demonstrated that hypoxia in the heart also increases mNCX activity (Hernansanz-Agustín et al., 2020).

The increase in mNCX activity and resistance to Ca^2+^-induced mPTP opening was not only observed in mitochondria isolated from CR hearts, but was also reproduced in cardiomyocytes incubated for 24 h with serum from CR animals (**Figure 3**), which exhibited increased NCLX and TMEM65 protein levels (**Figure 4**). These findings demonstrate that key mitochondrial adaptations induced by CR can be rapidly reproduced by exposure to circulating factors, consistent with previous observations that several CR phenotypes can be recapitulated by serum treatment in cultured cells (Cerqueira et al., 2016; de Cabo et al., 2003; Munhoz et al., 2023). This greatly broadens the translational potential of these mechanisms, suggesting that the benefits of CR may be achievable without prolonged dietary restriction itself.

Most importantly, the cardioprotective effects observed in both experimental models (cultured cardiomyocytes subjected to simulated I/R, **Figure 5**, and isolated perfused hearts exposed to I/R, **Figure 7**) were completely abolished by pharmacological inhibition of mNCX with CGP. This demonstrates that enhanced mitochondrial Na⁺/ Ca^2+^ exchange is not only associated with CR-induced cardioprotection, but is a mediator of its protective effects against ischemic injury. This conclusion is compatible with genetic studies showing that cardiac-specific deletion of NCLX results in mitochondrial Ca^2+^ overload, energetic collapse, and rapid heart failure and mortality (Luongo et al., 2017), while disruption of TMEM65 impairs mitochondrial Ca^2+^ extrusion and compromises cardiac function (Garbincius et al., 2025; Zhang et al., 2025). Conversely, increasing mitochondrial Ca^2+^ efflux by overexpression of NCLX has been shown to attenuate pathological cardiac remodelling and improve cardiac function in experimental heart failure (Garbincius et al., 2022). Collectively, these findings, together with our results, identify preservation of the TMEM65/NCLX complex and mNCX as a critical determinant of the cardioprotective phenotype induced by CR, and highlight enhancement of mitochondrial Ca^2+^ extrusion as a promising therapeutic strategy to limit ischemia/reperfusion injury.

## 5. CONCLUSION

CR protects the heart against IR injury by preserving mitochondrial Ca^2+^ homeostasis. This protection is associated with enhanced Na^+^-dependent mitochondrial Ca^2+^ efflux, resulting in improved mitochondrial function, reduced H_2_O_2_ release, delayed mPTP opening, and preserved cardiac performance after reperfusion. Pharmacological inhibition of mNCX abolished these benefits, demonstrating a central role for mitochondrial Ca^2+^ extrusion in CR-induced cardioprotection.

## ACKNOWLEDGEMENTS

This work was supported mainly by Fundação de Amparo à Pesquisa do Estado de São Paulo (FAPESP) grants 20/06970-5 and 13/07937-8, Conselho Nacional de Pesquisa e Desenvolvimento (CNPq), Coordenação de Aperfeiçoamento de Pessoal de Nível Superior (CAPES) line 01, Instituto Nacional de Ciência e Tecnologia em Metodologias Quantitativas e de Precisão em Biomedicina Redox grant #408213/2024-8 and the Centro de Pesquisa, Inovação e Difusão de Processos Redox em Biomedicina - CEPID Redoxoma grant 2013/07937-8. MICQ is supported by a FAPESP fellowship (24/07592-5). We thank Sirley M. de Oliveira and Camille C. C. da Silva for excellent technical assistance, Marcos V. Caetano, Manuel A. H. Lopes, and Lucas R. P. Camara for experimental help, as well as the expert animal facilities team lead by Silvania Neves and José Galeote Molero Leme de Oliveira. Artificial intelligence tools were used to improve language flow and organize references prior to proofreading by the authors, who are fully responsible for the content, data and text presented.

