## Supplementary material for "Caloric Restriction Promotes Ischemia/Reperfusion Cardioprotection Through Increased Mitochondrial Na^+^/Ca^2+^ Exchange": suplemental figures

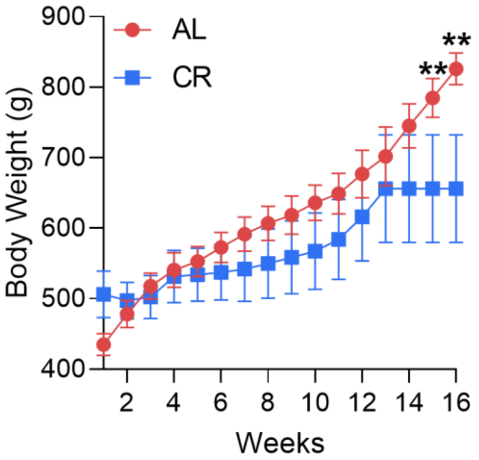


**S1. Effect of caloric restriction on body weight.** Body weight was monitored throughout the experimental period in animals from the *ad libitum* (AL) and caloric restriction (CR) groups. Data are presented as mean ± SD (n = 5). Statistical analysis was performed using t tests for each treatment week, ** $p < 0.01$.

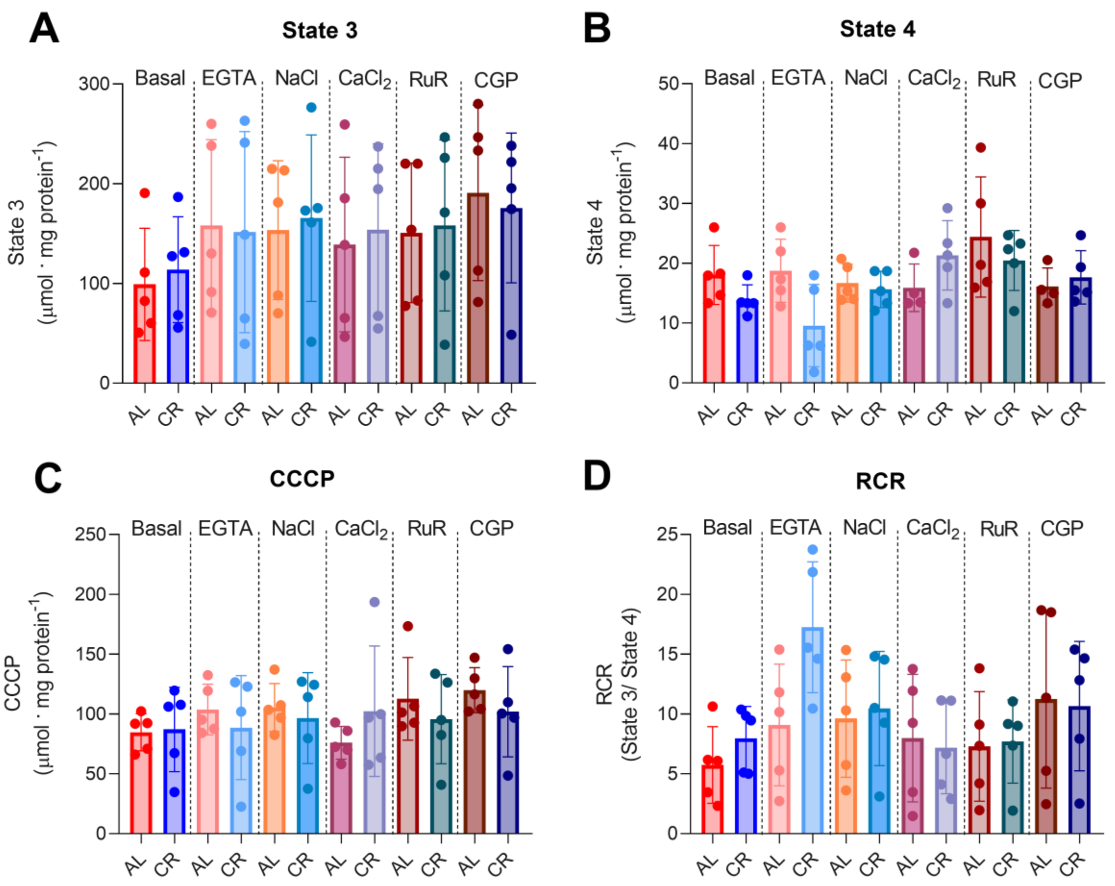


**Figure S2. Respiration of isolated heart mitochondria from control and calorie-restricted animals.** Heart mitochondrial oxygen consumption was assessed as described in Methods using 2 mM malate and 2 mM glutamate as substrates, followed by the addition of 1 mM ADP (State 3), 1 µM oligomycin (State 4), and 1.25 µM CCCP. The incubation medium contained 20 mM NaCl, 50 µM $CaCl_2$, 1.25 mM EGTA, 0.5 µM RuR, and 5 µM CGP, where indicated. (A) State 3, (B) State 4, (C) oxygen consumption in the presence of CCCP (State 3u), and (D) Respiratory Control Ratios (RCR, State 3/State 4). Data are presented as means ± SD (n = 5), and individual symbols represent biological replicates. Statistical analysis was performed using one-way ANOVA, with no significant differences.

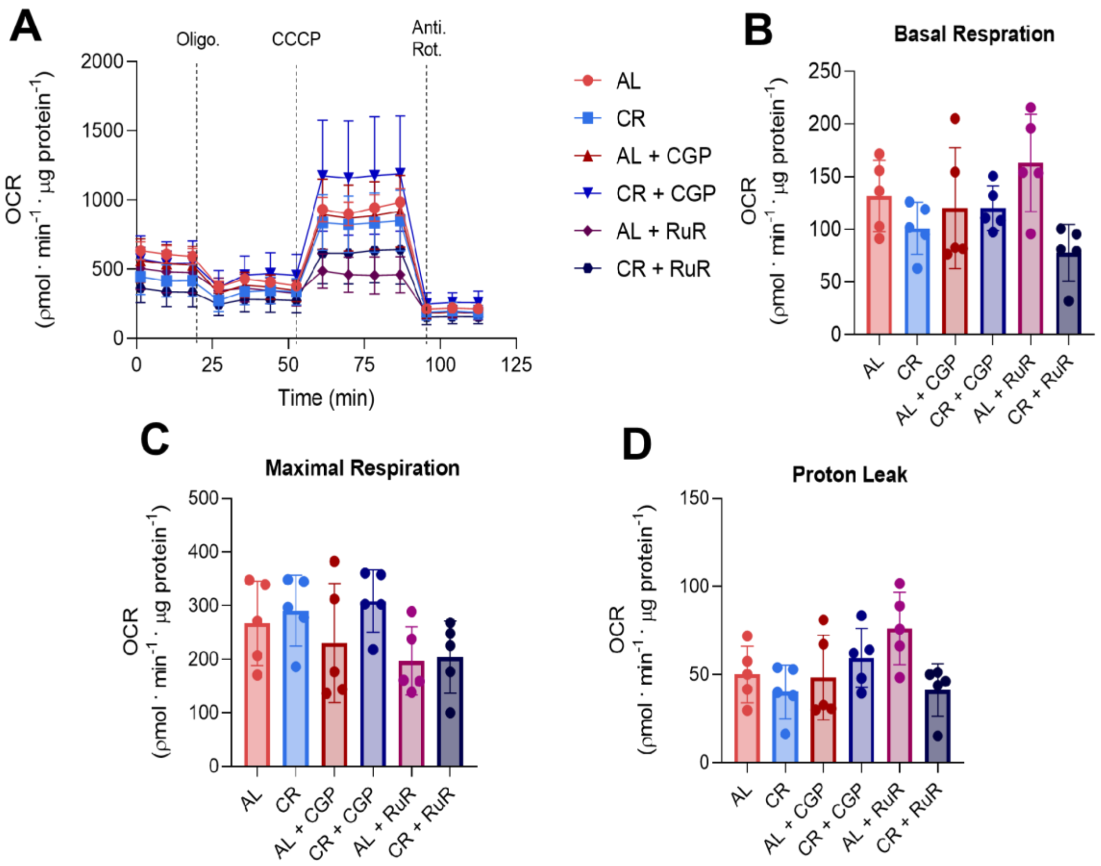


**Figure S3. Caloric restriction serum effects on oxygen consumption in cardiomyocytes.** (A) Representative oxygen consumption rate (OCR) traces of HL-1 cells treated with serum from *ad libitum*-fed (AL) and calorie-restricted (CR) animals, with or without treatment with RuR or CGP-37157, as indicated. (B) Basal OCR, (C) Maximal OCR (following CCCP addition), and (D) Proton leak (calculated as the difference between the minimum OCR after oligomycin addition and non-mitochondrial respiration). Data are presented as means ± SD (n = 5), and individual symbols represent biological replicates. Statistical analysis was performed using one-way ANOVA, with no significant differences.

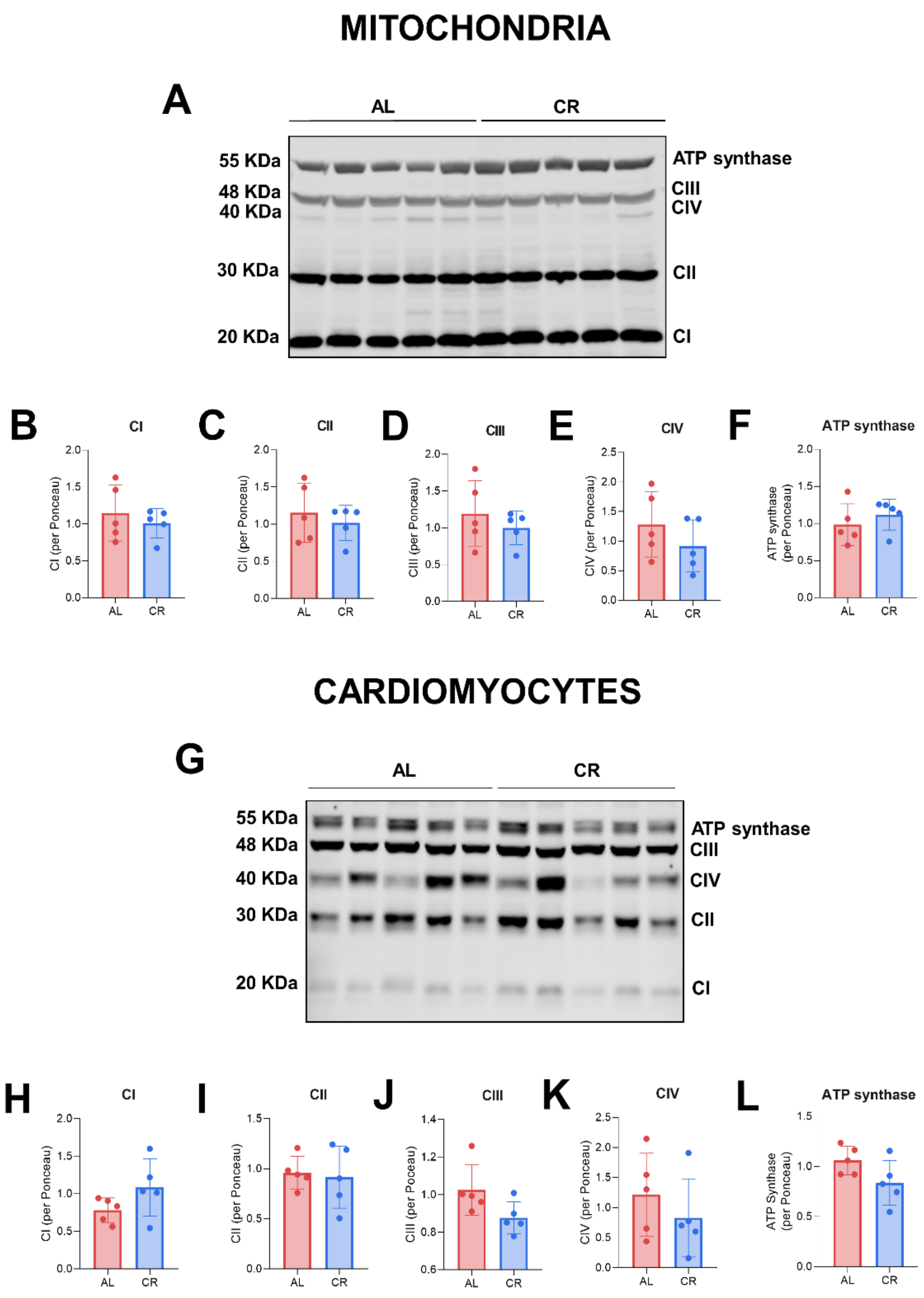


**Figure S4. Mitochondria isolated from CR animals and HL-1 cells treated with serum from CR animals show no differences in total OXPHOS protein levels compared to AL controls.** (A) Representative western blot (WB) of total OXPHOS proteins in isolated mitochondria from AL and CR animals, using 30 µg of total protein, (B–F) Quantification of the OXPHOS Western blot, showing protein levels of Complex I (CI), Complex II (CII), Complex III (CIII), Complex IV (CIV), and ATP synthase (CV), respectively. (G) Representative Western blot (WB) of total OXPHOS proteins in HL-1 cardiomyocytes treated for 24 h with serum from *ad libitum*-fed (AL) and calorie-restricted (CR) animals, using 30 µg of total protein, (H–L) quantification of the OXPHOS Western blot, showing protein levels of Complex I (CI), Complex II (CII), Complex III (CIII), Complex IV (CIV), and ATP synthase (CV), respectively. Data are presented as mean ± SD (n = 5), and individual symbols represent biological replicates. Statistical analysis was performed using t tests, with no significant differences.

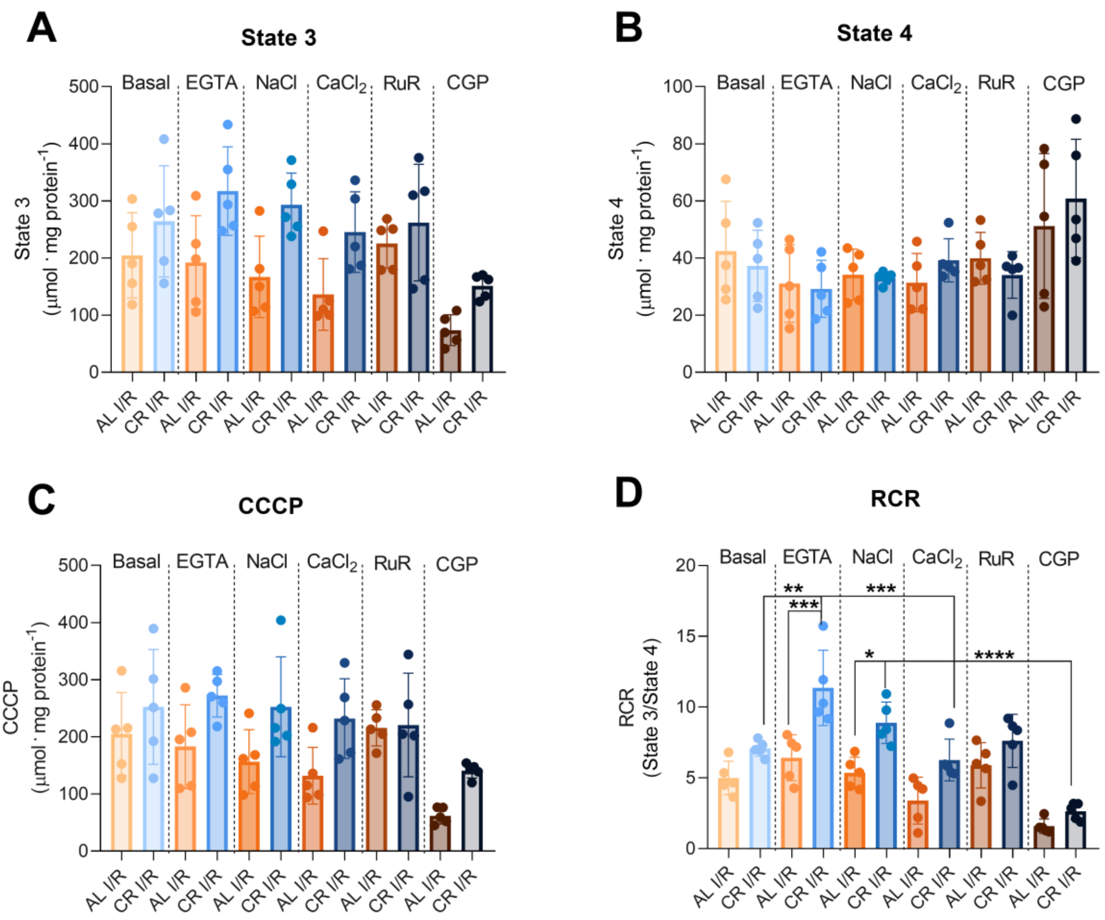


**Figure S5. Caloric restriction maintains respiratory control ratios after I/R.** Mitochondria were isolated after I/R, and respiration was assessed using 2 mM malate and 2 mM glutamate as substrates (Basal), followed by the addition of 1 mM ADP (State 3), 1 µM oligomycin (State 4), and 1.25 µM CCCP (CCCP). The incubation medium contained 20 mM NaCl, 50 µM $CaCl_2$, 1.25 mM EGTA, 0.5 µM RuR, and 5 µM CGP. (A) State 3, (B) State 4, (C) oxygen consumption in the presence of CCCP (State 3u), and (D) Respiratory Control Ratio (RCR, State 3/State 4). Data are presented as mean ± SD (n = 5). Statistical analysis was performed using one-way ANOVA, and individual symbols represent biological replicates. Statistical analysis was performed using one-way ANOVA, * $p < 0.05$; ** $p < 0.01$; *** $p < 0.005$.

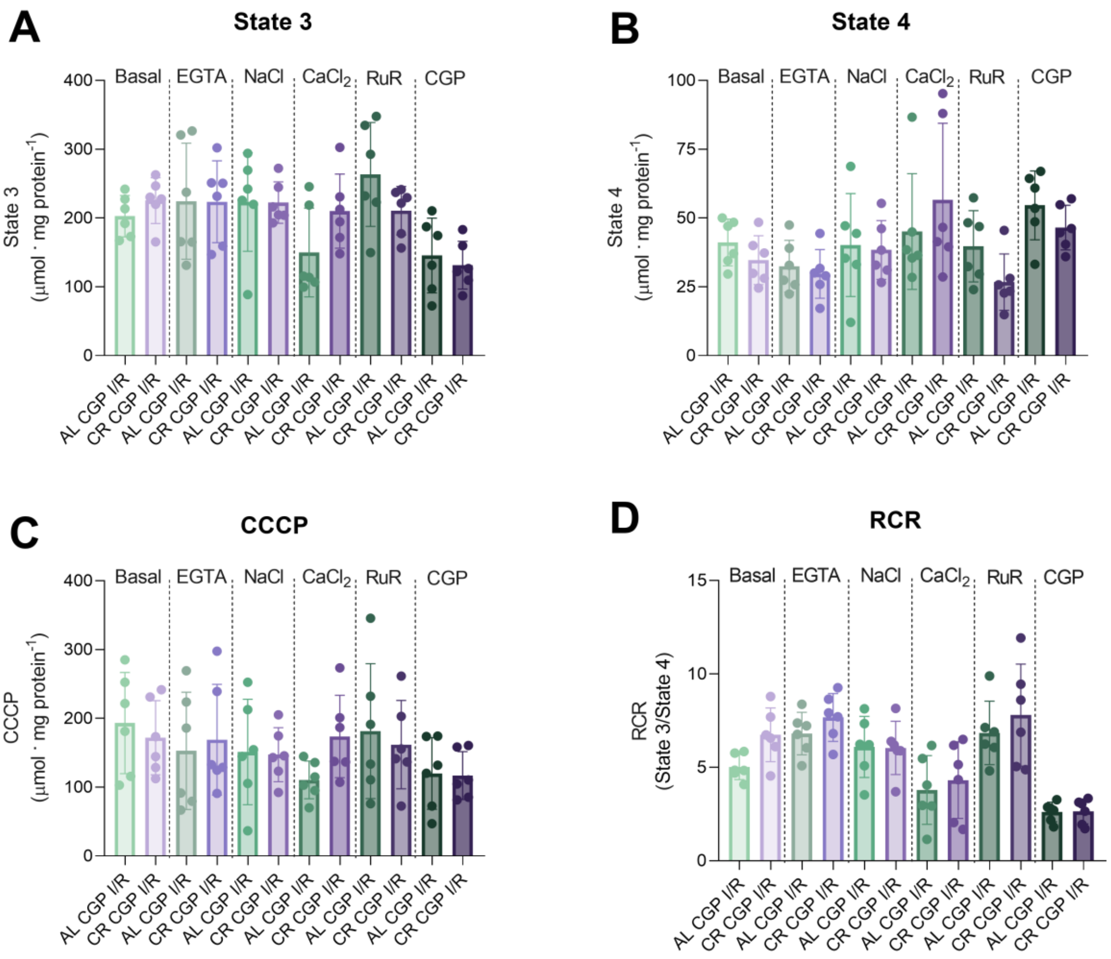


**Figure S6. The Na$^+$/Ca$^{2+}$ exchanger inhibitor CGP reverses the protection mediated by CR in respiratory control ratios.** Mitochondria were isolated after I/R in the presence of CPG, and respiration was assessed using 2 mM malate and 2 mM glutamate as substrates (Basal), followed by the addition of 1 mM ADP (State 3), 1 μM oligomycin (State 4), and 1.25 μM CCCP (CCCP). The incubation medium contained 20 mM NaCl, 50 μM $CaCl_2$, 1.25 mM EGTA, 0.5 μM RuR, and 5 μM CGP. (A) State 3, (B) State 4, (C) oxygen consumption in the presence of CCCP (State 3u), and (D) Respiratory Control Ratio (RCR, State 3/State 4). Data are presented as mean ± SD (n = 6), and individual symbols represent biological replicates. Statistical analysis was performed using one-way ANOVA, with no significant differences.

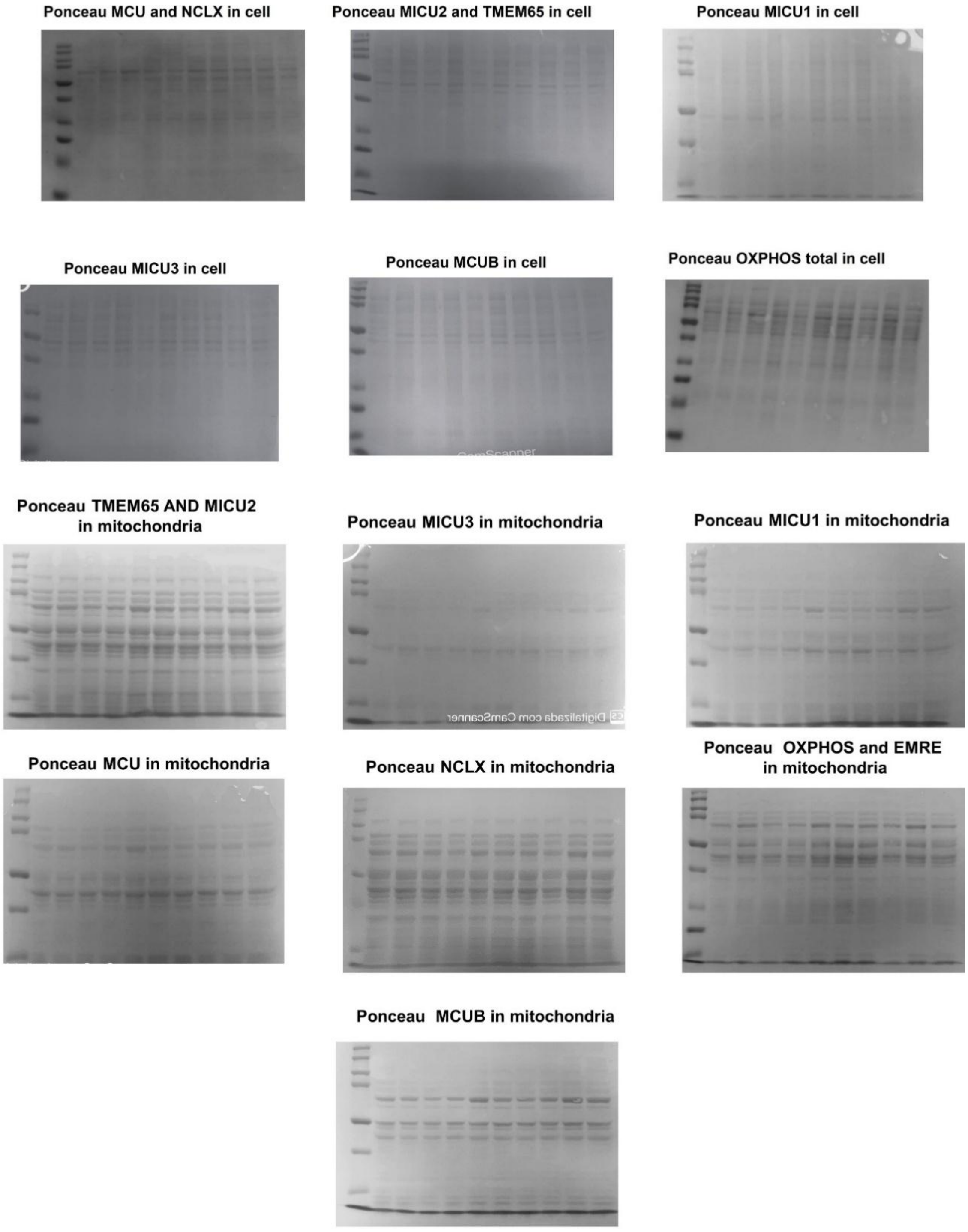


**Figure S7.** Ponceau used as western blot normalizers in cells and mitochondria. OxPhos data are presented in in Supplementary Figure 4, while all other loading controls correspond to blots presented in Figure 4. From upper left: MCU and NCLX in cells: Fig. 4K, 4P; MICU2 and TMEM65 in cells: Fig. 4M, 4Q; MICU1 in cells: Fig. 4L; MICU3 in cells: Fig. 4N; MCUB in cells: Fig. 4O; OXPHOS in cells: Fi. S4H-L; TMEM65 and MICU2 in mitochondria: Fig. 44D, 4I; MICU3 in mitochondria: Fig. 4E; MICU1 in mitochondria: Fig. 4C; MCU in mitochondria: Fig. 4B; NCLX in mitochondria: Fig. 4H; OXPHOS and EMRE in mitochondria: Figs. 4G and S4B-F; MCUB in mitochondria: Fig. 4F.

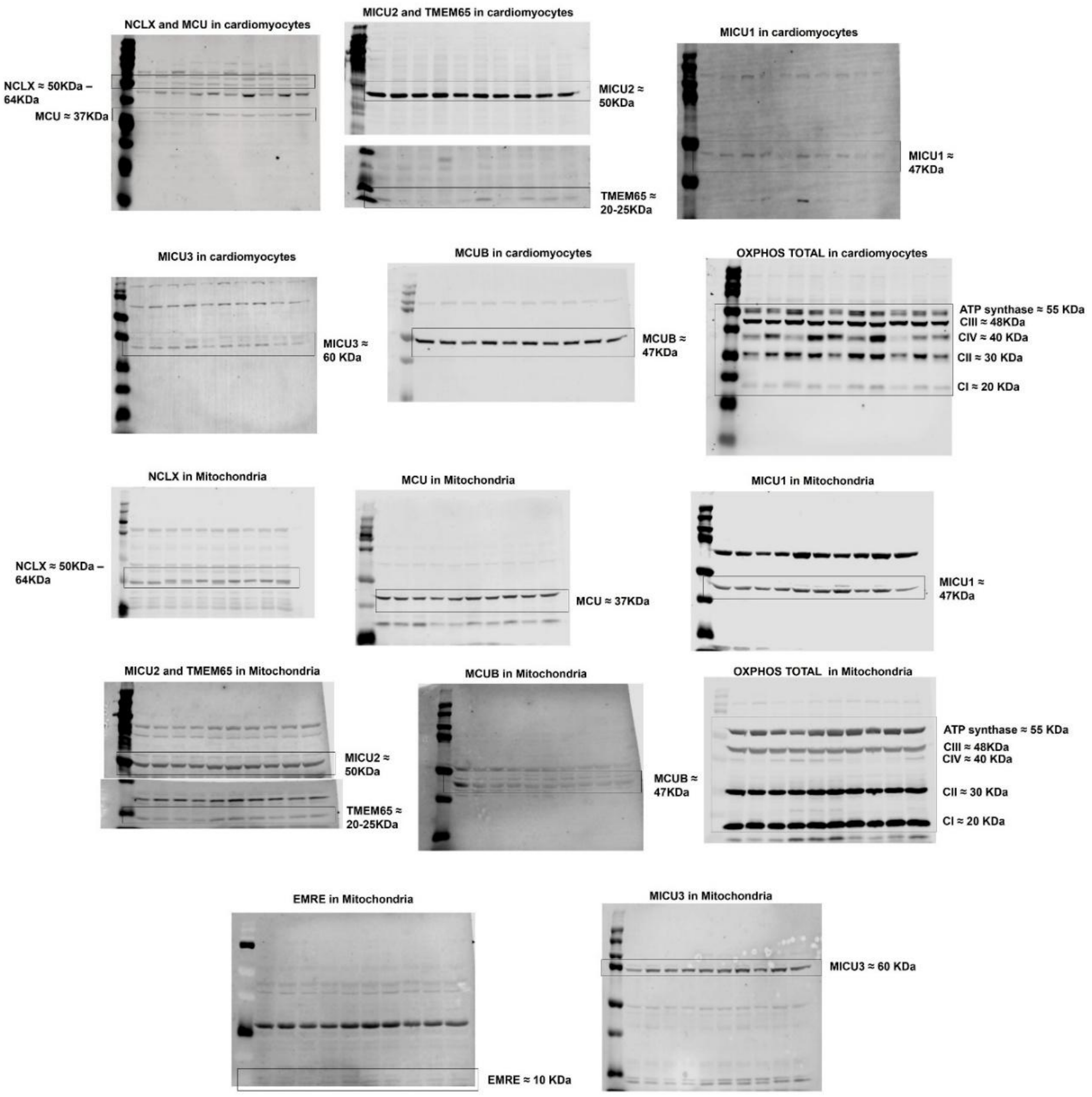


**Figure S8.** Full images of blots. Note membranes for MICU2 + TMEM65 were cut prior to staining for proteins in distinct mass ranges. Total OxPhos images are presented in cropped versions in Supplementary Figure 4, while all other images correspond to blots presented in cropped form in Figure 4. From upper left: NCLX and MCU in cardiomyocytes: Fig. 4J, 4K, and 4P; MICU2 and TMEM65 in cardiomyocytes: Fig. 4J, 4M, and 4Q; MICU1 in cardiomyocytes: Fig. 4J and 4L; MICU3 in cardiomyocytes: Fig. 4J and 4N; MCUB in cardiomyocytes: Fig. 4J and 4O; OXPHOS in cardiomyocytes: Fig. S4G-L; NCLX in mitochondria: Fig. 4A and 4H; MCU in mitochondria: Fig. 4A and 4B; MICU1 in mitochondria: Fig. 4A and 4C; MICU2 and TMEM65 in mitochondria: Fig. 4A, 4D, and 4I; MCUB in mitochondria: Fig. 4F; OXPHOS in mitochondria: Fig. S4A-F; EMRE in mitochondria: Fig. 4A and 4G; MICU3 in mitochondria: Fig. 4A and 4E.
